# Synthesizing images to map neural networks to the human brain

**DOI:** 10.1101/2025.11.25.689428

**Authors:** Kailong Peng, Kenneth A. Norman, Nicholas B. Turk-Browne, Jeffrey D. Wammes

**Affiliations:** Department of Psychology, Yale University, New Haven, CT, USA; Interdepartmental Neuroscience Program, Yale University, New Haven, CT, USA; Department of Psychology, Princeton University, Princeton, NJ, USA; Princeton Neuroscience Institute, Princeton University, Princeton, NJ, USA; Wu Tsai Institute, Yale University, New Haven, CT, USA; Department of Psychology, Queen’s University, Kingston, ON, Canada; Centre for Neuroscience Studies, Queen’s University, Kingston, ON, Canada

**Keywords:** Image Synthesis, Convolutional Neural Network (CNN), fMRI, Representational Similarity Matrix (RSM), Residual Method, Brain Mapping, Visual Cortex

## Abstract

Computational models can be used to generate hypotheses about the brain. In the visual system, this approach has revealed similarities between how natural images are represented in convolutional neural networks and in cortical regions. However, natural images generate highly correlated representations across the hierarchy of model layers, meaning that each brain region will not correspond selectively to a single layer. Because each model layer performs additional image transformations, gaining a more selective mapping between individual layers and regions could reveal how specific algorithmic stages of processing are instantiated in the brain. To enable this mapping, we developed a generative framework for image synthesis that minimizes the similarity of image representational similarity matrices across model layers, aiming to orthogonalize the distances between the patterns of unit activations evoked by the same set of images. With the patterns orthogonalized, the resulting similarity matrix for each layer provides a fingerprint of the unique computational role of that layer. To test this approach, we synthesized 16 artificial images from the Inception-V1/GoogLeNet model and scanned participants with fMRI while they viewed these images repeatedly in random order. Image-specific patterns of voxel activity were used to compute image-by-image similarity matrices across the whole brain with searchlights. Most layers could be mapped to circumscribed cortical regions, and these mappings overlapped less than the mappings obtained with natural images. Given the prevalence of existing fMRI datasets with natural images, we used the synthesis method as a benchmark to develop an alternative residual method that can achieve comparable performance for natural image datasets. These approaches could be extended to other neural network architectures and stimulus modalities for targeted mappings of model computations to the brain and behavior.

## Introduction

Advances in computer vision have enabled rapid progress in testing theories about the organization and function of the visual system. Mapping from activation in the units of neural networks — which can be easily engineered, trained, and tested — to activation in the brain has improved our understanding of the nature of computations in visual cortex. Findings from these studies have established parallels between computer vision models and the brain at various levels of abstraction (1–3). When models are trained to produce human-like performance, their units predict neuronal responses in primate visual cortex (4, 5). Through careful tailoring of architectures and training sets, both supervised (6–9) and unsupervised neural networks (10, 11), have been able to capture properties of biological vision.

Scaling up to the level of populations of neurons, as captured by non-invasive human brain imaging methods like functional magnetic resonance imaging (fMRI), the similarity of activation patterns across images matches the similarity of unit patterns in models (3, 12–14). Further, lower layers in the model tend to predict earlier areas along the ventral visual stream, while higher model layers predict later areas (2, 15), suggesting that these models may have a brain-like processing hierarchy. Representations in the hidden layers of these networks are sensitive to shape (16), animacy (17), object-scene relationships (12), center-peripheral spatial dynamics (18), and other visual properties. These models therefore establish a useful analogy to how the human visual system is instantiated in the brain, reinforced by the finding that model representations can predict human behavioral judgments (19).

However, a major challenge in using models to understand computations in the brain is that the mapping between model layers and visual areas is imprecise. A given model layer can be reflected in multiple visual areas, and each visual area can be predicted from multiple model layers. These blurred lines and the distributed mapping of model layers in the brain may be real, but an alternative possibility is that it results from the fact that natural images produce highly correlated representational structures across adjacent layers of a convolutional neural network. This leads to a many-to-many mapping, preventing the identification of more specific links between model layers and visual areas. To address this challenge, we developed a model-based method of creating novel abstract image sets that can distinguish the unique representations encoded at multiple stages of visual processing in the brain.

Optimizing image sets for the expression of specific model features is a form of image synthesis, which was initially developed to provide a window into what neural networks ‘see’ or have learned (20, 21). However, image synthesis has since evolved into a critical tool for investigating the parallels between neural network features and brain functions. Given the correspondence of low to high model layers with early to late ventral stream regions, respectively (2, 15), image synthesis can be used to target the representational organization of visual areas. For example, to investigate higher-order visual cortex, image sets have been synthesized that equate features in lower layers of a neural network while manipulating the correspondences between patterns of unit activity in higher layers, allowing for control over representations in later regions in the ventral visual processing stream (22).

Here we extend this concept and target multiple areas of visual cortex simultaneously by manipulating feature relationships across multiple layers of a convolutional neural network. The goal of our method is to synthesize a single image set that exhibits a unique representational structure in *every* model layer, allowing for brain responses evoked by these images to be mapped more precisely to individual model layers. We then showed the synthesized images to human participants while their brains were scanned using fMRI. To anticipate the results, our synthesis approach revealed a more selective mapping of model layers to cortical regions than was possible with natural images. Because of the widespread availabilty of valuable fMRI datasets with natural images, we also present a second method based on residual regression that can be used with such datasets to achieve similarly precise mappings.

## Materials and methods

### Image synthesis

We developed an approach to create image sets that allow unique mappings from individual layers of a convolutional neural network (CNN) to the human brain. The main objective for such image sets is that the similarity of unit activation patterns for all pairs of images — the representational similarity matrix (RSM) — in each layer is orthogonal to the RSM of all other model layers. This criterion is not met with natural images. To generate images, we used the Inception-V1 model architecture (also referred to as GoogLeNet; Depth/Blocks: 22; Layers: 144) (23).

There are three main reasons why we selected a CNN model in general and this specific architecture in particular. First, in pilot studies, Inception-V1 generated the most perceptually diverse and interesting images, making it suitable for psychological experiments with humans in the fMRI scanner. Second, in our prior work, we were able to generate images that parametrically manipulated pairwise image similarity in higher-order visual cortex (22). Third, several prior studies have mapped CNN layers to brain regions in the visual processing hierarchy (3). We do not claim that this is the best model for object or visual recognition in the brain; other models may offer superior performance. However, CNNs have been widely studied in terms of their correspondence with brain activity and often result in many-to-many mappings between model layers and cortical regions. The aim of this paper is to explore whether the same types of models, used generatively, could lead to a more selective mapping. In Inception-V1, we focused on the first three convolutional layers and all subsequent mixed-pooling layers. In addition to achieving the desired feature correspondences across layers, we also aimed to ensure that the images were visually rich and distinguishable. To this end, we started with 16 visual noise arrays (Fig. 1A), which were optimized iteratively 28,800 times to enhance the activation of one of the learned feature channels from the model. We ensured that the images contained a broad distribution of RGB values, promoting images that were more vibrant and saturated in color.

**Figure 1.**
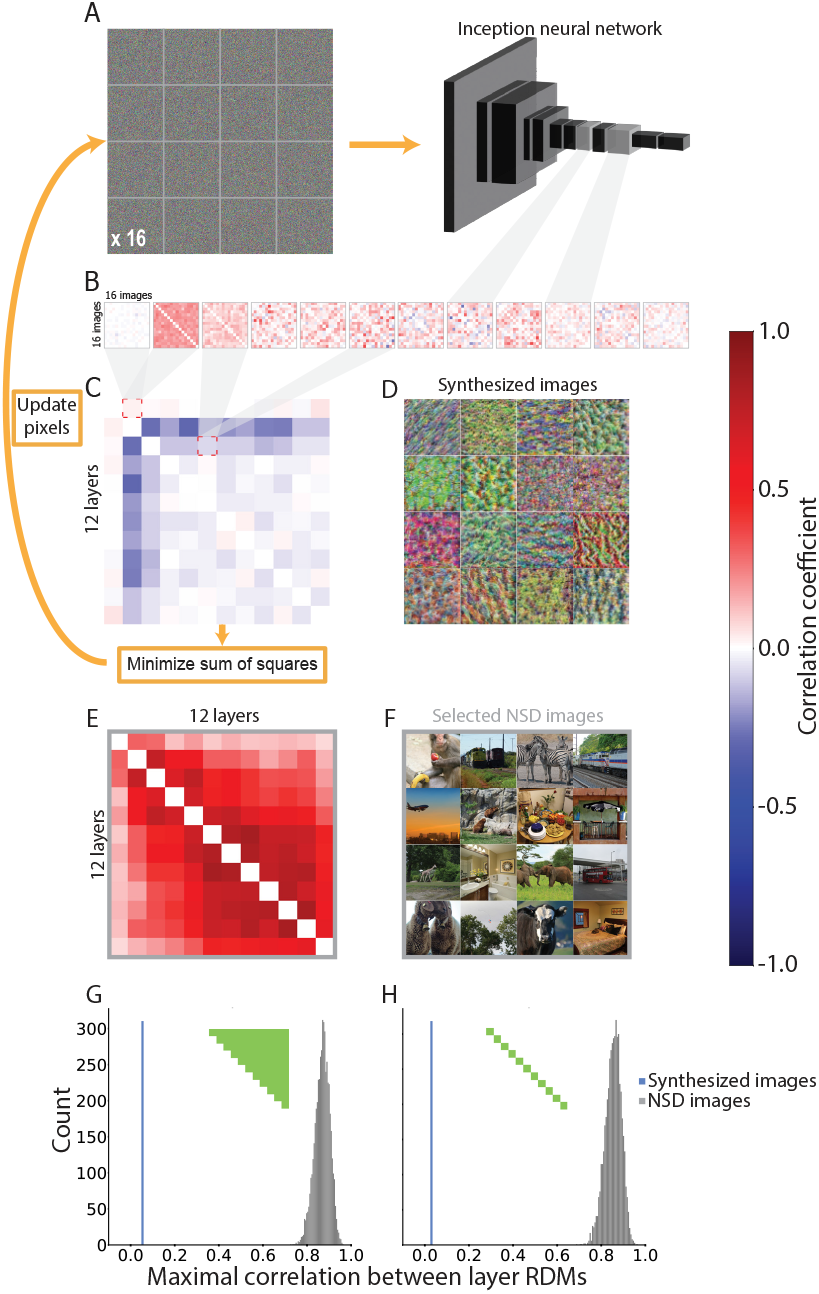
Image synthesis process. (A) The 16 synthesized images begin as 16 random visual noise images input to the CNN. (B) The pattern of activation for each image across units in a given model layer is then correlated with the pattern for every other image to generate a 16 image by 16 image RSM for each layer. (C) These 12 RSMs are vectorized and correlated with each other to create a 12 layer by 12 layer second-order correlation matrix. This matrix is squared and summed, providing a single metric that captures the overall similarity of representational structures across model layers. This metric is used as the cost function to update pixels in the synthesized images through backpropagation such that the sum of squares is iteratively minimized. (D) The final synthesized images successfully elicited a second-order correlation matrix with minimal values; that is, RSMs for each of 12 model layers were as distinct as possible from one another. (E) As a control, we computed the second-order correlation matrix for sets of 16 natural images randomly selected from Natural Scenes Dataset (NSD). The RSMs for these images were more strongly correlated across layers than the RSMs for the synthesized images. (F) One of the natural image sets from NSD. (G) Average correlation between all pairs of layers (green portion of inset RSM) for the final synthesized images (blue line) was lower than any of the average layer correlations from 5,000 sets of 16 natural images from NSD (gray histogram). (H) The result was virtually identical when focusing on the average second-order correlations of adjacent layers.

Using these 16 colorful images as a starting point, we computed the 16 image x 16 image RSM describing the relationships (i.e., Pearson correlations) between all pairs of unit activation patterns for the 16 images in each of the 12 selected layers (Fig. 1B). This resulted in 12 layer-specific RSMs. Next, we computed the second-order similarity between all pairs of these layer-specific RSMs, producing a 12 layer x 12 layer correlation matrix describing the relationships among the representational structure in each layer (Fig. 1C). Finally, we calculated the sum of squares of this correlation matrix, providing a single metric that, when minimized, enhances the distinctiveness of the representational structures across layers. On each of 1,800 iterations, the pixels of the 16 images were simultaneously updated with the optimization objective of reducing this second-order sum of squares metric using gradient descent.

### fMRI task design

We collected up to eight task runs (375 s each) of fMRI data from 30 human participants. During each run, participants viewed five repetitions of each of the 16 synthesized images. The images were presented one at a time in the center of the screen (Fig. S1). The sequence of images in each run was constructed pseudo-randomly, with the constraint that each of the 16 synthesized images would be presented exactly 5 times each (for a total of 80 image presentations) without back-to-back repetitions. Each image was presented for 1.5 s, with inter-stimulus intervals between trials jittered with equal probability at 1.5, 3, or 4.5 s. To ensure attention, participants were required to monitor the images for infrequent trials (10% of trials) where a small square within the displayed image was presented in grayscale instead of its normal color. Participants indicated that they detected a target image by pressing a button on a handheld button box.

### Participants

Thirty healthy young adults (age range: 19–33 years, 19 female) with normal or corrected-to-normal visual acuity took part in the fMRI experiment. All participants provided written informed consent before the experiment and received payment for their participation. The experiments were approved by the Institutional Review Board at Yale University. The experiment was completed in a single session lasting for 1.5 hours.

### Data acquisition

Data were acquired using a 3T Siemens Prisma MRI scanner with a 64-channel head coil in the Brain Imaging Center at Yale University. Eight functional runs were collected from most participants (*n*=23), but delays or other constraints led to seven (*n*=4) or six (*n*=3) runs in some participants. Functional data were acquired using a multiband echo-planar imaging (EPI) sequence (TR = 1,500 ms; TE = 32 ms; voxel size = 1.5 mm isotropic; flip angle = 64°; multiband factor = 6), yielding 90 axial slices. Each run contained 250 volumes. For field-map correction, two spin-echo field map volumes (TR = 11,220 ms; TE = 66 ms) were acquired in opposite phase-encoding directions; the field maps otherwise matched the parameters of our functional acquisitions. We also collected a whole-brain T1-weighted magnetization prepared rapid gradient echo (MPRAGE) image (TR = 1,960 ms; TE = 2.26 ms; voxel size = 1 mm isotropic; flip angle = 8°; GRAPPA acceleration factor = 2; 208 sagittal slices), a high-resolution T2-weighted turbo spin-echo (TSE) image (TR = 11,170 ms; TE = 93 ms; voxel size = 0.4 × 0.4 × 1.5 mm; flip angle = 150°; distance factor = 20%; 54 coronal slices perpendicular to the long axis of the hippocampus), and a whole-brain T2-weighted turbo spin-echo (TSE) image (TR = 3,200 ms; TE = 565 ms; voxel size = 1 × 1 × 1 mm; flip angle = Variable; 176 sagittal slices).

We also performed a secondary analysis on the Natural Scenes Dataset (NSD), which consists of 7T fMRI data from eight participants viewing scene images (age range: 19–32 years, 6 female). See the original paper for full details on the task and sequences (24).

### Data preprocessing

We preprocessed each functional run using the FEAT tool in the FMRIB Software Library (FSL) (25). Preprocessing included brain extraction, slice-timing correction, high-pass filtering with a 100-s cutoff, and alignment to the run’s middle functional volume using MCFLIRT (26). We also used FSL’s topup tool (27) to address susceptibility-induced distortions with the field maps. The functional runs were aligned to each participant’s T1-weighted anatomical image through boundary-based registration and to the standard MNI space using FLIRT (26, 28), with 6 and 12 degrees of freedom, respectively. Analyses of individual runs were performed in the participant’s native EPI space. Group-level analyses were conducted in standard MNI space.

### Defining image representations

We analyzed functional runs using FSL’s FEAT tool (25, 29, 30). We generated an experimental design file for each of the 16 images by convolving the onsets for that image with a double-gamma hemodynamic response function (HRF). This produced a regressor for each of the 16 images in each run and participant. With this design, we used a mass-univariate general linear model (GLM) to estimate every voxel’s average BOLD response to each image. The GLM therefore produced voxelwise wholebrain maps of beta weights for each image in each run, reflecting the characteristic pattern of brain activity (i.e., the representation) for that image. These maps were transformed into the participant’s T1-weighted anatomical space and standard MNI space.

### Searchlight analysis of representational stability

We conducted a whole-brain searchlight analysis to validate these image representations by testing their stability across runs. Surrounding each voxel in the whole-brain map, we extracted the patterns of activity from a 7 × 7 × 7 cube of voxels. We tested whether this pattern for one image and run was more similar to the pattern for that same image in another run than it was to the patterns for other images in the other run (Fig. S2). For each participant and searchlight, we computed within-versus between-image similarity across runs for all 16 images (A, B, C, …, P). Within-image similarity was calculated as the average Pearson correlation between presentations of the same image across all pairings of the eight runs. For example, image A in run 1 was correlated with image A in run 2, run 3, run 4, etc.; image B in run 1 was correlated with image B in run 2, run 3, run 4, etc. This led to 448 total comparisons (8 choose 2 for each of the 16 images).

Between-image similarity was calculated as the average Pearson correlation between searchlight patterns of activity for different images across all pairings of the eight runs. For example, image A in run 1 was correlated with image B, image C, etc. up to image P, in run 2, run 3, run 4, etc. Because this produced a much larger number of pairings than the within-image similarity metric (8 choose 2 × 16 × 15 = 6,720) we randomly subsampled 448 values to match the number of values averaged into the estimate of within-image similarity. This resampling procedure was repeated 5,000 times to generate a sampling distribution of between-image similarity values. To quantify the representational stability in each searchlight, we computed the z-score of the within-image similarity with respect to the distribution of the resampled between-image similarities. This metric served as the output value for each searchlight and was computed across the whole brain for each participant, yielding a voxelwise map of representational stability. The statistical significance of the stability at each searchlight was calculated at a group level in standard space with FSL’s randomise and corrected for multiple comparisons with threshold-free cluster enhancement (TFCE). The resulting map of stable representations was used to generate masks for later analyses.

For the secondary NSD dataset, 16 sets of 16 natural images were chosen. Following the same procedure as for the synthesized images, in each participant and image set we conducted a searchlight analysis by computing within-versus between-image neural similarity throughout the brain. These searchlight results were then averaged across the 16 sets within-participant and statistical significance was assessed at the group level with randomise and TFCE.

### Searchlight analysis of model layers with synthesized images

After establishing stability, our main analyses focus on establishing model-to-brain mappings. To achieve this, we used a whole-brain searchlight approach to localize model layers within the brain at the group level. We first compiled the fMRI data in standard space for all participants. Next, within each searchlight, we calculated a 16 image by 16 image RSM by correlating the beta values from the voxels within the searchlight for all pairs of the 16 synthesized images. We then averaged these RSMs across all but one held-out participant and fit a linear regression model predicting the upper triangle of the averaged matrix for the searchlight from the corresponding upper triangle of the 16 image by 16 image RSM describing the relationships among unit activation patterns in a selected model layer. The learned model-to-brain regression weights were then used to predict the held-out participant’s RSM. This prediction was evaluated against the actual searchlight RSM from the held-out participant with R-squared. We cross-validated this analysis such that each participant was held out and the whole procedure was repeated for each of the 12 model layers, resulting in a cross-validated R-squared statistic for each layer in each searchlight.

We normalized these R-squared values against an empirical null distribution established by shuffling the upper triangle of each searchlight’s RSM, thereby obliterating any real associations between model and brain that might have been present. In these scrambled data, the entire leave-one-participant-out cross-validated linear regression analysis detailed above was completed, and this process, with unique sets of scrambled data, was repeated 500 times. The true R-squared for each searchlight was converted to a z-score with respect to this null distribution. The result was a whole-brain, voxelwise searchlight map of z-scores for each layer and participant reflecting how well the representational space of that model layer explained the representational space of their local brain responses. The statistical significance of these model-layer fits was calculated at the group level for each layer with randomise and TFCE. For the mapping between model layers and neural representations to be meaningful, a prerequisite is that a given cluster of voxels shows stable image representations. Therefore, we restricted analyses to the voxels established in the preceding analysis that exhibited representational stability across runs.

### Searchlight analysis of model layers with NSD images

The searchlight analysis pipeline above for the synthesized images was repeated for each of the 16 sets of 16 NSD images. The z-score searchlight maps for each layer were averaged across image set within-participant prior to group analysis. We also analyzed each set separately at a group level to check the consistency across sets.

### Searchlight analysis of model layers with NSD images with residual method

A critical extension of the above analyses — the “residual method” — was designed to identify brain patterns uniquely associated with each model layer even when using NSD images, which normally overlap considerably across model layers. The core objective was to remove all variance in each searchlight RSM that could be explained by all model-layer RSMs *other than* the model layer currently being fit. To achieve this, we added a residualizing step in the linear regression model fitting. The resulting fit of the RSM for that model layer to the residual of the searchlight RSM thus reflects the unique variance the model layer explains with respect to the other model layers. This provides an alternative to our image synthesis approach that can allow the identification of orthogonalized brain mappings for each model layer in the absence of image synthesis (i.e., using preexisting natural images).

The residual method was performed for a model layer of interest by predicting the upper triangle of the searchlight RSM with a multiple linear regression containing 11 predictors corresponding to the upper triangles of all other layers; the upper triangle of each model layer’s RSM and each searchlight’s RSM were z-scored in advance. The residual of this regression (i.e., the unexplained variance in the searchlight RSM) was extracted and fit with a simple linear regression that used the upper triangle of the held-out model layer’s RSM as its sole predictor. Otherwise, the remainder of the pipeline mirrored that for the synthesized images.

### Overlap analysis

To assess the degree of neural overlap between the searchlight map for one base layer (M) with respect to another layer (N), we computed the ratio of the number of significant voxels (after correction) shared between the two maps (intersection of M and N) to the total number of significant voxels in the base map (M). In other words, what proportion of the map M was shared with N. This resulted in a 12 layer by 12 layer overlap matrix. Note that this matrix can be asymmetric, as the proportion of M shared with N does not necessarily equal the proportion of N shared with M. For example, consider what would happen if M is a subset of N with only half as many significant voxels: the ratio M:N would be 1.0, whereas the ratio N:M would be only 0.5.

To assess whether the image synthesis method improved the specificity of the mappings between model layers and the brain, we compared the mean of all off-diagonal elements of the overlap matrix for synthesized images to the distribution of mean overlaps of the searchlight maps from the NSD images. We used bootstrap resampling to generate this distribution. After computing a separate overlap matrix and the off-diagonal mean for each of the 16 NSD image sets, we randomly sampled 16 values from these 16 means with replacement and calculated the grand mean of these 16 values. This was repeated 5,000 times to produce a distribution with which the mean overlap for the synthesized images could be compared. We repeated this same analysis with a focus on the overlap of only adjacent model layers (elements one off the diagonal, both above and below), as opposed to all possible pairings of layers.

As mentioned, we also developed a residual regression method to test whether it would allow for layer-specific mappings to be found even with natural images. To determine whether this method was as effective as our approach, we also compared the mean overlap for our synthesized images to the distribution that results from applying the residual method to the NSD images.

## Results

### Image synthesis

We used image synthesis to generate an artificial stimulus set with distinct representational similarity patterns across layers of a CNN model. The algorithm began with a set of 16 randomly initialized images and iteratively refined the images using backpropagation, with the objective of minimizing the similarity of RSMs across all 12 layers. This procedure was sucessful in reducing the pairwise correlations of layer-specific RSMs to around zero (Fig. 1C). This was considerably lower than for natural images (Fig. 1E; *p ≪* 0.0001), both for all pairs of model layers (Fig. 1G) and when only considering adjacent model layers (Fig. 1H). We therefore used the synthesized image set (Fig. 1D) for the fMRI task.

### Representational stability

The synthesized images were shown to participants repeatedly during fMRI. Because they were novel and abstract stimuli, we first asked which brain regions exhibit stable representations of the images across repetitions, to provide a basis for later mapping of model layers. We used a searchlight analysis to assess the stability of the neural representations of the images. Specifically, we identified clusters of voxels that exhibited higher across-run similarity for repetitions of the same image compared to presentations of two different images (TFCE corrected, *p <* 0.05). We observed representational stability throughout visual cortex in the occipital and parietal lobes, extending to regions of lateral prefrontal cortex (Fig. S3A). This widespread representational stability indicates that our synthesized images elicited reliable patterns of neural activity throughout the visual processing hierarchy, ranging from lower-order to higher-order regions. This cortical map of representational stability was converted to a binary mask, which was applied to select which voxels were used in subsequent analyses. This ensured that we were only attempting to use the model to predict neural representations that were themselves reliable.

A parallel analysis was performed for natural images (averaging across 16 NSD image sets). The representations of natural images were likewise most stable in visual cortex, extending into ventral temporal cortex (Fig. S3B).

### Mapping model layers in the brain

We used a searchlight analysis to identify which brain regions contained representations that mirrored the representations encoded in each layer of the CNN model. The logic of using synthesized images is that the representational structures of these images were orthogonalized across layers (by design), with the goal of obtaining more selective layer mappings in the brain. We used a leave-one-participant-out linear regression approach to obtain these mappings. For each model layer and searchlight, a linear regression was fit between the layer’s RSM for the synthesized images and the brain’s RSM for the synthesized images, using all participants except for one held out participant. The regression model’s performance was tested by attempting to predict the held-out participant’s RSM in that searchlight. This procedure was repeated across all searchlights for each participant, resulting in a group-level map of the expression of each of the 12 selected layers (TFCE corrected, *p <* 0.05). We then used the results from our previous representational stability analysis to generate a mask so that the layer mappings to the brain included only areas that had consistent image representations. Model layers 1-5, 7, 9-10, and 12 were significantly represented in the brain (Fig. 3).

**Figure 2.**
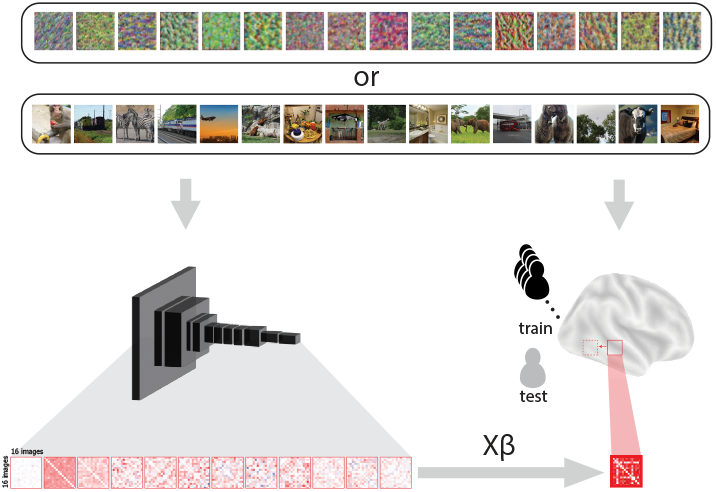
Mapping model layers to the brain. Sets of 16 synthesized images and 16 natural images were presented to both human participants and a CNN model. Activity patterns for each of the 16 images in a set were extracted across voxels in a local brain searchlight and across all of the units in a selected model layer. We computed the pairwise correlation of these activity patterns across the images in each set, resulting in a 16 image by 16 image RSM for the brain searchlight and the model layer. For each model layer independently, we fit a leave-one-participant-out cross-validated linear regression model predicting the searchlight RSM from the layer RSM with data from the entire sample (‘train’; black) except for one held-out participant (‘test’; gray). We finally tested the regression model using the learned weights (*Xβ*) to predict the searchlight RSM of the held-out participant, and evaluated the prediction against their actual searchlight RSM. This produced a measure for each participant of how well every model layer explained the representational structure of each searchlight in the brain, which we normalized to the variance explained by a randomized null distribution.

**Figure 3.**
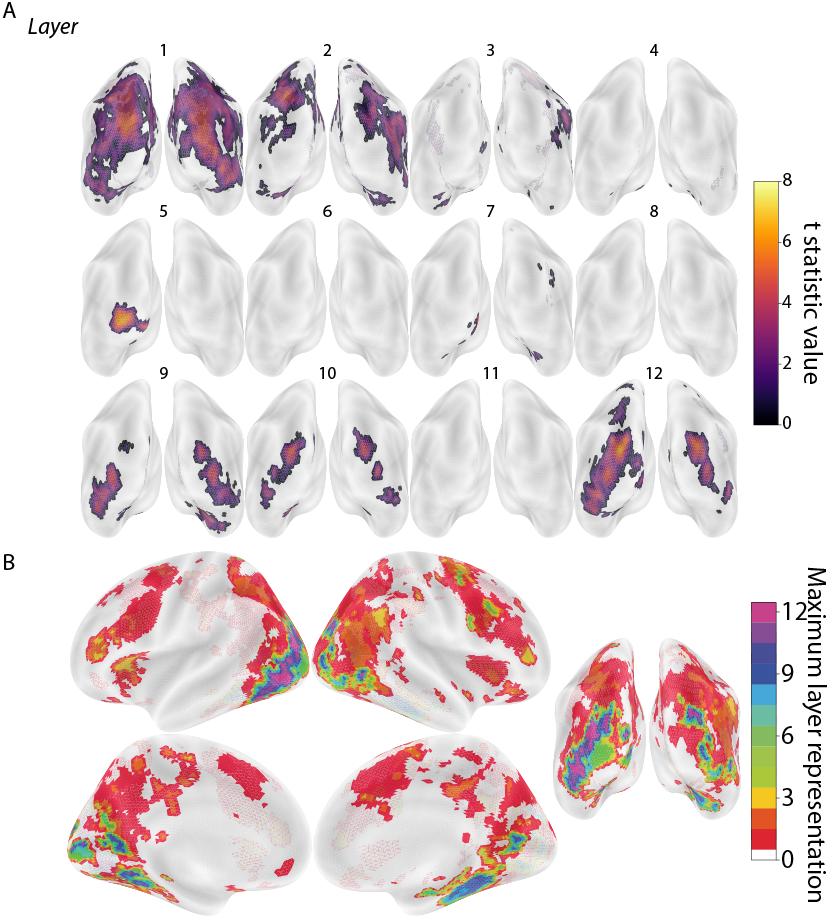
Localization of model layers based on synthesized images. Leave-one-participant-out linear regression analysis predicting the RSMs of fMRI activity patterns in each searchlight from RSMs of unit activation patterns in the 12 CNN model layers for the synthesized images. (A) Posterior view of whole-brain searchlight results for each model layer masked based on representational stability. Depicted clusters survived whole-brain correction for multiple comparisons (TFCE, *p <* 0.05). The large clusters for layer 1’s mapping may reflect weak but broadly distributed signals that are retained as significant by the TFCE method; in support of this, a more traditional cluster-thresholding approach yielded a more selective map (Fig. S4). (B) Consolidated map of all layers with voxels labeled according to which (if any) model layer had the maximum t-statistic. For example, if the t-statistic for layer 12 exceeded that of all other layers in a region, the region was colored purple.

Model layers were also localized in the brain using 16 sets of 16 natural images from NSD. The searchlight results for each layer were first averaged across the 16 sets within-participant, resulting in a group-level map for the 12 selected layers, analogous to the synthesized images (TFCE corrected, *p <* 0.05). However, in contrast to the synthesized images, the representations of natural images in the CNN layers were expressed broadly throughout occipital, parietal, and temporal lobes and, critically, in a very similar way across all layers (Fig. 4). That is, natural images appear to produce considerably more overlap across layers than the synthesized images, which we quantify in the next section.

**Figure 4.**
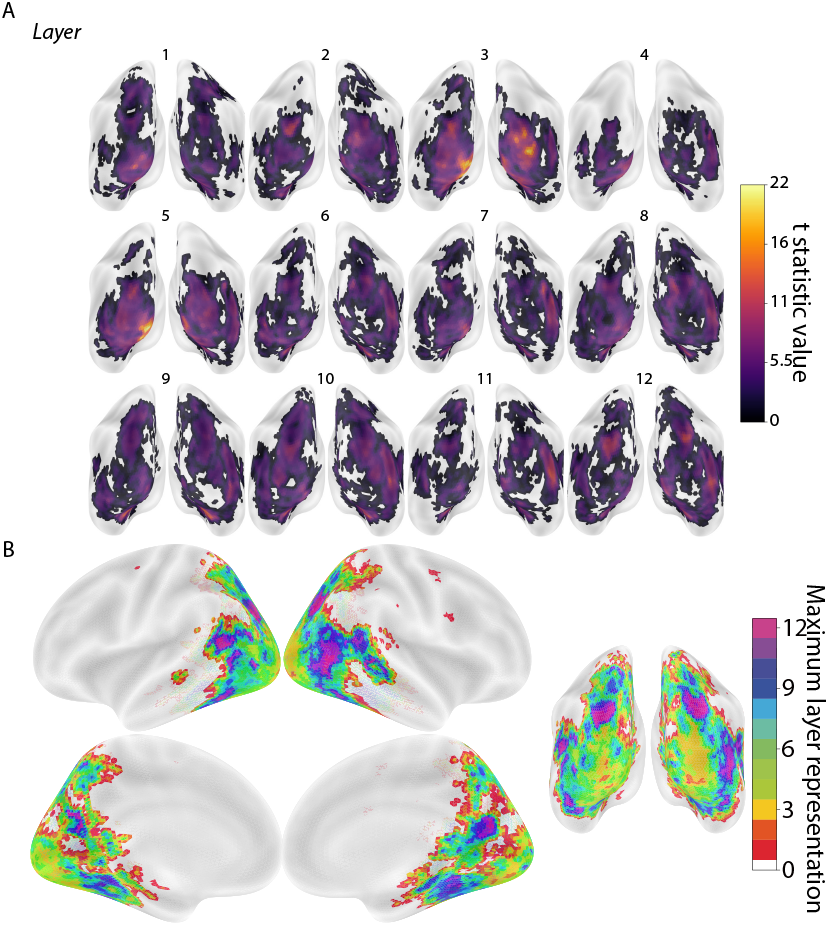
Localization of model layers based on natural images. Leave-one-participant-out linear regression analysis predicting the RSMs of fMRI activity patterns in each searchlight from RSMs of unit activation patterns in the 12 CNN model layers for the natural images. The results have been averaged over 16 sets of natural images from NSD, though there was some variability across sets (Fig. S5). Posterior view of average whole-brain searchlight results for each model layer masked based on representational stability. Depicted clusters survived whole-brain correction for multiple comparisons (TFCE, *p <* 0.05). (B) Consolidated map of all layers with voxels labeled according to which (if any) model layer had the maximum t-statistic.

### Overlap analysis

We hypothesized that synthesizing images with the objective of orthogonalizing the representational similarity of those images across layers would lead to a more selective mapping of the layers in the human brain. We quantified this selectivity by computing the normalized overlap of the searchlight maps for different CNN layers with synthesized vs. natural images. The overlap of the map for one layer (M) with respect to another layer (N) was calculated as the ratio of voxels that were significant (TFCE corrected, *p <* 0.05) in both M and N to the total number of voxels that were significant in M (Fig. 5A).

**Figure 5.**
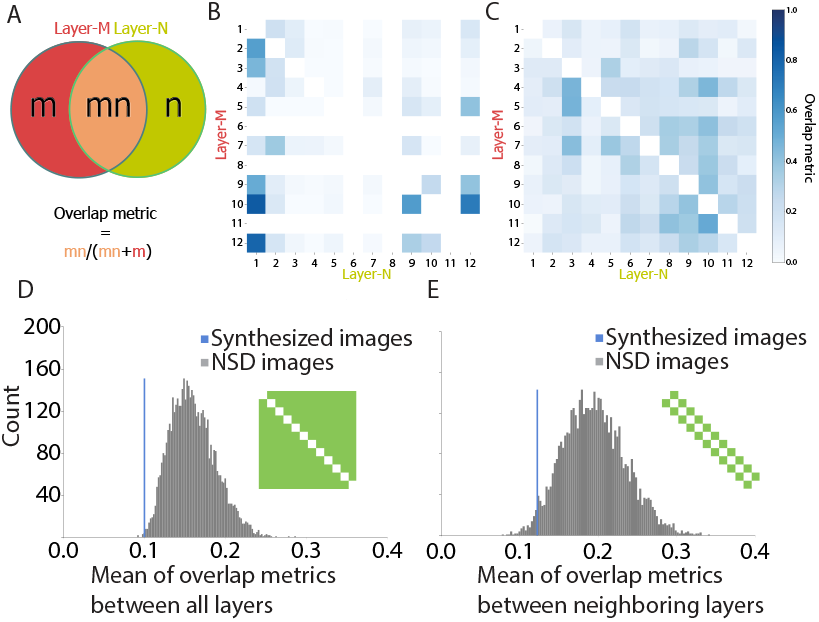
Overlap of searchlight maps for synthesized and natural images. (A) Venn diagram with the two circles (red and green) indicating the set of significant voxels in the searchlight maps for two layers (M and N): *m* is the voxel subset unique to M, *n* is the voxel subset unique to N, and *mn* is the voxel subset shared between M and N. The overlap of M with N was defined as the ratio of shared voxels (*mn*) divided by the total number of M voxels (*mn* + *m*); likewise, the overlap of N with M was shared (*mn*) divided y total N voxels (*mn* + *n*). (B) The overlap metrics between all pairs of 12 layers in both directions for the synthesized images. (C) The overlap metrics between all pairs of 12 layers in both directions for the natural images from NSD. (D) We used bootstrap resampling to compare the average pair-wise overlap metric for searchlight maps generated from synthesized vs. natural images. We estimated the overlap for synthesized images (blue line) as the mean of off-diagonal values (excluding layers 6, 8, and 11 with no significant voxels) in the layer overlap matrix from subpanel B. This mean was compared to a sampling distribution of the off-diagonal mean overlaps for natural images (gray histogram) derived by bootstrap resampling of the 16 NSD image sets with replacement 5,000 times. (E) The same procedure as subpanel D restricted to adjacent model layers.

This overlap analysis revealed a robust difference in the degree of overlap between the mappings of CNN layers for synthesized vs. natural images. Specifically, the synthesized images produced less overlap for almost all layer comparisons (Fig. 5B) than natural images from NSD (Fig. 5C). This observation was verified statistically by comparing the mean off-diagonal layer overlap for the synthesized images to a sampling distribution for natural images estimated with bootstrap resampling across image sets (*p* = 0.0018; Fig. 5D). This result replicated when focusing on only adjacent layers most likely to overlap (*p* = 0.0036; Fig. 5E). These findings suggest that image synthesis helps localize and differentiate brain regions preferentially involved in layer-specific computations.

### Residual method

Although image synthesis may be an efficient way to find selective representations of model layers in the human brain, this requires the collection of new fMRI data, which comes at the cost of time and resources. We thus developed an alternate “residual method” that can achieve similar layer selectivity with existing fMRI datasets in which participants view natural images. Rather than synthesizing stimuli to orthogonalize RSMs across model layers, we instead used a regression approach to orthogonalize the RSMs obtained with natural images prior to final analysis, isolating the variance in representational structure unique to a given layer. Namely, using multiple linear regression for each layer and searchlight, we fit a model predicting the RSM of fMRI activity in each searchlight with the RSMs from all model layers *except for* one held-out layer. The residuals of this model represent the unique variance in the representational structure of that searchlight that remains unexplained by the other 11 layers. We then used the RSM from the held-out 12th layer as a predictor to fit to this remaining residual variance. The amount of layer-unique variance was quantified in a leave-one-participant-out manner.

The mapping of individual model layers to the brain (TFCE corrected, *p <* 0.05) was more selective with this residual method than in the original analysis of natural images, in terms of the searchlight maps (Fig. 6A-B) and layer overlap matrix (Fig. 6C). Using the residual method, the amount of layer overlap in the brain for natural images was not statistically different than for synthesized images both across all layer pairs (*p* = 0.85; Fig. 6D) and when restricted to adjacent layers (*p* = 0.96; Fig. 6E). This demonstrates that the residual method achieved comparable selectivity to image synthesis while leveraging an existing natural image dataset.

**Figure 6.**
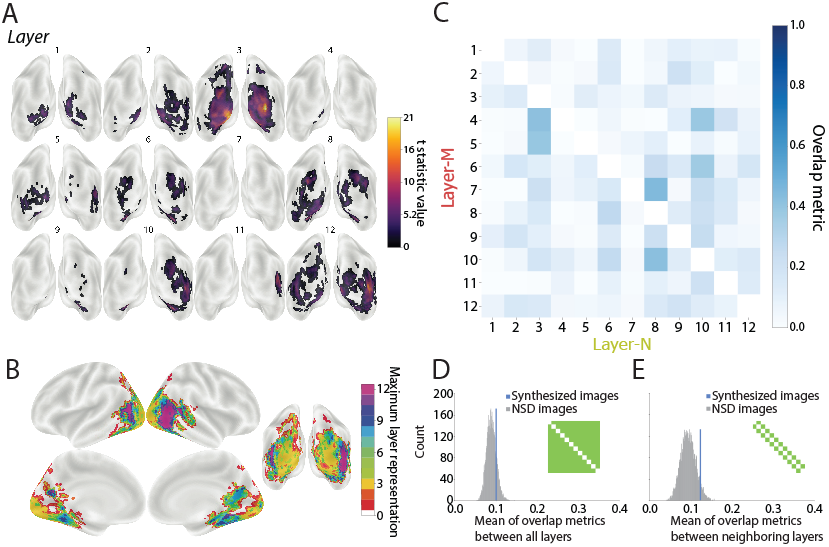
Localization of model layers based on natural images with residual method. Leave-one-participant-out linear regression analysis that predicted RSMs throughout the brain from the RSM for one model layer after regressing out all other model layers. The results have been averaged over the 16 sets of natural images from NSD, though there was variability in localization across sets (Fig. S6). (A) Posterior view of average whole-brain searchlight results for each model layer. Depicted clusters survived whole-brain correction for multiple comparisons (TFCE, *p <* 0.05). (B) Consolidated map of all layers with voxels labeled according to which (if any) model layer had the maximum t-statistic. (C) The overlap metrics between all pairs of 12 layers in both directions for the natural images from NSD after the residual method. (D) Comparison of mean off-diagonal of layer overlap matrix for synthesized images (blue line) to the bootstrapped sampling distribution for natural images after residual method. (E) The same procedure as D but restricted to overlap metrics from adjacent layers.

## Discussion

We developed a synthesis method for generating artificial image sets that are almost entirely distinct in their representational geometry across the layers of a CNN model. This enabled layer-specific predictions that mapped selectively to the brain. Strikingly, the majority of model layers (1–5, 7, 9–10, 12) could be mapped to cortical regions that were both anatomically consistent and generalizable across participants. Our synthesis approach also yielded much lower overlap across layers than what was obtained with sets of natural images.

This study builds on previous evidence that neural networks provide a useful analogy to the brain (2, 15, 22). Following the lineage of work that has engineered single images to manipulate visual-evoked activity in macaque (1) and human brain regions (31), our method, using a relatively small set of images, can drive unique *relationships* between patterns of activity across several brain regions simultaneously (c.f. (22)). Critically, this synthesis method increases experimental efficiency by allowing studies to target specific visual features or stages of processing with modest amounts of data.

Although we found layer-selective cortical regions for many of the model layers using synthesized images, their localization did not follow an obvious hierarchical structure in the brain, which appears inconsistent with related work (2, 15). This difference could be related to the dependence of CNN models on low-level features and textures (see Fig. 2 of (32)), which are traditionally associated with early visual cortex (33, 34). This existing tendency may be further compounded by the abstract nature of the synthesized images, which do not contain the identifiable discrete objects, scenes, or compositional structure that would traditionally be the domain of higher-level or category-selective visual areas like LO and IT. Future studies will be needed to synthesize images based on features from later layers or from other model architectures, including those that have been trained to be less dependent on texture (32).

However, it is not always feasible to collect a new fMRI dataset with synthesized images, and there are many available datasets with naturalistic images (24, 35, 36). To identify precise layer representations in the brain using such existing datasets, we developed a residual method that attained comparable performance to the synthesized images in terms of distinctive layers. Further, unlike the synthesized images, residualized naturalistic images revealed a pronounced hierarchical structure (Fig. 4B and 6B). While this is desirable, the model-to-brain correspondences were inconsistent (Fig. S6), generating variable layer fingerprints. This highlights the inherent limitations of using existing data: mapping stability is constrained by the scale and quality of the dataset. Thus, while residual methods are well-suited for exploratory research with limited resources, synthetic approaches offer superior flexibility and precision for hypothesis-driven studies.

The high correlation across model layers for natural images raises the question of how the synthesis method successfully orthogonalized RSMs. One interpretation is that the synthesized images occupy parts of the representational landscape where feature transformations are most prominent across layers (see Supporting Information; Fig. S7). By enforcing good coverage of this part of the space, synthesis captures the unique transformation of each layer with relatively few stimuli, something that the manual selection of natural images cannot guarantee. Though synthesis may highlight layer-specific transformations, both synthesized and residualized natural images led to partial overlap across model layers in the brain, highlighting the challenge of one-to-one layer-to brain region mapping. This may arise because the simplicity of feedforward CNNs, which do not reflect the complex additional recurrent connections that characterize the ventral visual stream (3, 37) and have been incorporated into newer models (e.g., (38)).

In sum, our findings indicate that computer vision models can be used generatively to develop stimulus sets capable of targeting and distinguishing different stages of visual computation in the brain. As a complement to this novel approach, we also developed a statistical technique for producing similar orthogonalization using pre-existing natural image datasets. Together, these approaches provide a toolkit that can be used for precise model-based interrogation of the complex computations that allow the human brain to achieve higher-order vision, which in turn, will inform the refinement of computational models as better analogs for the brain.

## Supporting information

Supplemental info part 1

Supplemental info part 2

## Code and data availability

The code and data of this study will be made publicly available following the acceptance of this manuscript.

## Acknowledgments

We thank the Turk-Browne Lab and the Yale Interdepartmental Neuroscience Program for support and feedback.

## Author Contributions

**Conceptualization:** Kailong Peng, Jeffrey D. Wammes, Kenneth A. Norman, Nicholas B. Turk-Browne;

**Methodology:** Kailong Peng, Jeffrey D. Wammes, Nicholas B. Turk-Browne;

**Data Collection:** Kailong Peng, Jeffrey D. Wammes;

**Writing - Original Draft Preparation:** Kailong Peng, Jeffrey D. Wammes;

**Writing - Review & Editing:** Jeffrey D. Wammes, Nicholas

B. Turk-Browne, Kenneth A. Norman;

**Supervision:** Nicholas B. Turk-Browne;

**Funding Acquisition:** Kenneth A. Norman, Nicholas B. Turk-Browne;

All authors have read and agreed to the published version of the manuscript.

