## Supplemental info part 1 for "Synthesizing images to map neural networks to the human brain"

### Supporting Information

#### Toy dataset to interpret the image synthesis process.

Here we adopt a manifold interpretation of neural networks (1, 2) to provide a crude intuition of how image synthesis might have worked in our study and its difference from the traditional mapping of models to the brain with randomly selected natural images. For a given set of images, neighboring layers in a CNN model should have similar representational distances. However, we were able to generate images that occupy specific points in representational space whose distances were close to orthogonal across neighboring layers. Crucially, this was accomplished without modifying the model weights. Through our image synthesis process, we may have found pairs of points that capture the most substantial variations in the manifold between model layers. These variations play a crucial role in the manifold transformation, effectively defining the meaningful computations carried out within the network.

To provide a visual intuition (Fig. S7), we simulated clouds of data points in the shape of an S (layer 1) and a rectangle (layer 2). Each point in a given cloud represents a high-dimensional pattern of unit activations and has a corresponding point in the other cloud. Each cloud corresponds to the representational space of a model layer. From these clouds, 64 points were randomly selected and visualized (red) (Fig. S7AB). A process similar to image synthesis was applied to these 64 points, resulting in their relocation (Fig. S7CD). This process is designed to be analogous to the image synthesis method which is trying to make fingerprints of different model layers dinstinctive from each other. Here we are algorithmically optimizing the distance matrices between different dots(images) of layer 1 and layer 2 to be distinctive from each other. More specifically, the objective of this algorithm was to identify a set of points that elicited distinct RSMs within these manifolds. The RSM was defined as the pairwise distance of the 64 points, yielding 64 x 64 RSMs for layers 1 and 2. During the synthesis process, each iteration updated one of the 64 points. The 11 points nearest to the updating point (including itself) were identified. The updating point was sequentially replaced with each of these 11 points, resulting in a new configuration of 64 points. The new configuration was evaluated to assess if it better satisfied the optimization objective — to minimize the squared correlation between the RSMs of the two model layers. The point from the 11 candidates that best fulfilled the optimization objective was selected to replace the updating point. This iteration process continued until the optimization objective no longer decreased. As a result, a set of 64 points that achieved distinctiveness in the RSMs was identified. Notably, these points occupy manifold regions that undergo the most drastic changes across layers. For example, the yellow points in layer 1 (Fig. S7C) were closer to the teal than green points, but this reversed in layer 2 (Fig. S7D). In other words, the image synthesis algorithm identifies points that lie close to the S-manifold curves of layer 1 while being distant in the planar manifold of layer 2.


1. Pratik Prabhanjan Brahma, Dapeng Wu, and Yiyuan She. Why deep learning works: A


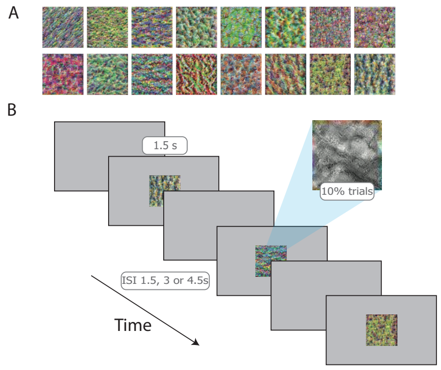


**Figure S1. Stimuli and scanner behavioral task.** (A) Set of 16 synthesized images, optimized to produce a unique RSM for each layer in a neural network model. (B) In each run, images were displayed for 1.5 s, followed by an inter-stimulus interval (ISI) jittered between 1.5, 3, and 4.5 s. To ensure attention, participants performed a cover task of pressing a button when they spotted a rare (10% of trials) small grayscale patch on an image.

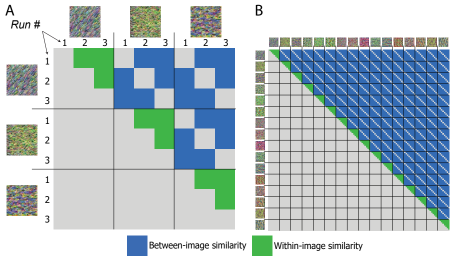


**Figure S2. Searchlight analysis of representational stability.** (A) Simplified schematic diagram depicting a hypothetical design matrix for three images across three runs. Within-image neural similarity across runs is contrasted with between-image neural similarity across runs. We used a searchlight analysis to find regions in the brain with greater within- versus between-image similarity. (B) The full design matrix used in the analysis, with 16 images across 8 runs.

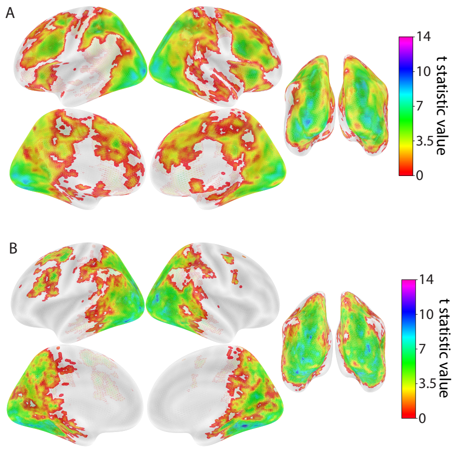


**Figure S3. Representational stability mask.** (A) A searchlight analysis was used to identify brain regions with greater within- than between-image similarity in neural responses for the synthesized images. Colors depict $t$-statistic for clusters that survived TFCE multiple comparisons correction ($p < 0.05$). Subsequent analyses mapping model layers to brain searchlights were restricted to this mask, as it would not make sense to ask how layers account for representations that are themselves unstable. (B) This same analysis was repeated for the 16 sets of 16 natural NSD images. We averaged the searchlight result across NSD sets.

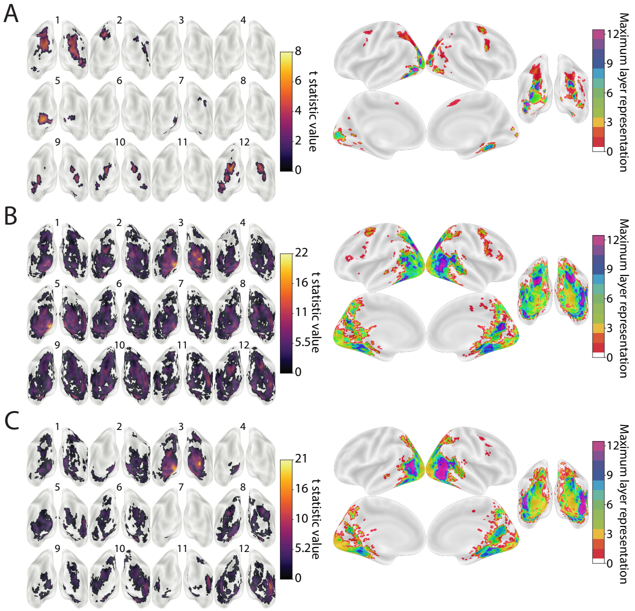


**Figure S4. Localization of model layers with cluster-mass correction.** The results of searchlight analyses localizing model layers in the brain using cluster-mass correction for multiple comparisons (cluster-forming threshold $p < 0.001$) rather than TFCE, for (A) synthesized images, (B) non-residual NSD images, and (C) residual NSD images. The broad map of layer 1 for the synthesized images in the original analysis (Fig. 3) was much reduced, suggesting that TFCE was capturing weak but highly distributed signals.

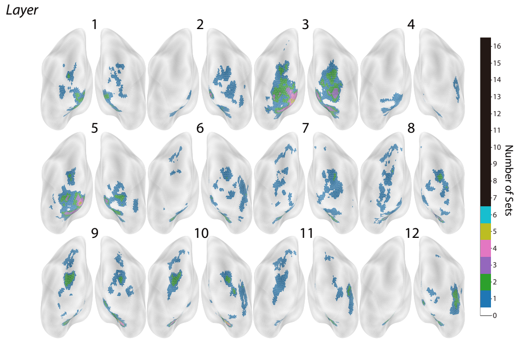


**Figure S5. Variation across NSD image sets for non-residual method.** We ran the full searchlight and group analysis separately for each of the 16 NSD image sets and then summed across the resulting clusters. The number of sets associated with each searchlight (which could range from 0–16) was relatively low (up to 6), suggesting that the mapping of model layers varied across image sets (see also Fig. S8).

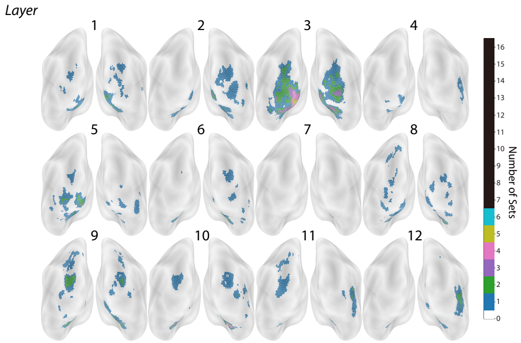


**Figure S6. Variation across NSD image sets for residual method.** We repeated the same analysis as in Fig. S5, but used the residual method. This again yielded low overlap across sets, suggesting that the mapping of model layers varied across image sets (see also Fig. S8).

manifold disentanglement perspective. *IEEE transactions on neural networks and learning systems*, 27(10):1997–2008, 2015.

2. Štefan Pócoš, Iveta Bečková, Tomáš Kuzma, and Igor Farkaš. Assessment of manifold unfolding in trained deep neural network classifiers. In *Trustworthy AI-Integrating Learning, Optimization and Reasoning: First International Workshop, TAILOR 2020, Virtual Event, September 4–5, 2020, Revised Selected Papers 1*, pages 93–103. Springer, 2021.
