## Supplemental info part 2 for "Synthesizing images to map neural networks to the human brain"

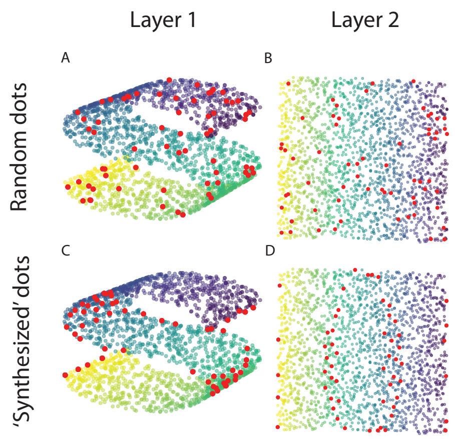


**Figure S7. Manifold visualization of image synthesis across two layers for simulated toy dataset.** (A) 64 points (red) were randomly selected from the S-shape representational manifold in layer 1. (b) The same 64 points are visualized in the unrolled representational manifold for layer 2. (c) After an optimization procedure similar to our image synthesis method, the 64 points are visualized in the representational manifold for layer 1. (d) The same points are visualized in the representational manifold for layer 2. The image synthesis objective shifts the points to parts of the manifold that undergo the most dramatic changes across layers.

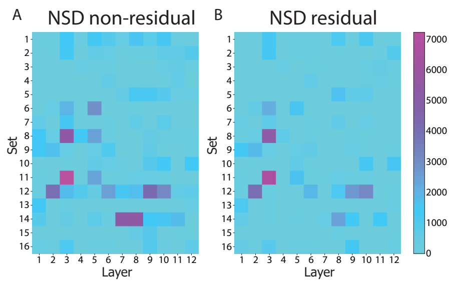


**Figure S8. Voxel count for each set of NSD images using non-residual and residual methods.** The number of significant voxels for each set of NSD images was determined for all 12 model layers. This information was obtained by conducting separate searchlight analyses for each of the 16 NSD sets. The expression of model layers was variable across NSD sets. These findings indicate that randomly selecting a small number of NSD images leads to instability in the mapping between the brain and the model.
